# Rare habitats, rare species, and invasive predators highlight management complexities in the Colorado River system

**DOI:** 10.1101/2024.12.15.628570

**Authors:** Blake R. Hossack, Kenzi M. Stemp, Caren S. Goldberg, Alexandra C. K. Duke, Taryn N. Preston, Jeff L. Arnold, Andrew M. Ray

## Abstract

Long-term drought caused Lake Powell, a reservoir on the Colorado River (USA), to decline to its lowest elevation in >50 years during 2022–2023, allowing warm water to pass through intakes of Glen Canyon Dam and facilitating invasion by non-native Smallmouth Bass (*Micropterus dolomieu*). Establishment of bass downstream of the dam could threaten persistence of several native fishes, including two federally listed species. Subsequent detection of larval Smallmouth Bass in a spring-fed slough (river mile -12 slough) connected to the river in Glen Canyon National Recreation Area (NRA) increased urgency to stem further invasion. The National Park Service is evaluating proposed actions to limit effects from non-native predators on native species in the Colorado River, including potentially channelizing the slough. This locally rare, spring-fed waterbody provides habitat for other species, including Western Tiger Salamanders (*Ambystoma mavortium* subsp.) of uncertain origin. We found salamanders from the slough had two distinct mitochondrial DNA haplotypes identical to sequences from nearby Arizona Tiger Salamander (*A. m. nebulosum*) populations, confirming they are the native genotype. We detected Red-spotted Toads (*Anaxyrus punctatus*) and Woodhouse’s Toads (*A. woodhousii*) from three other sites in Glen Canyon NRA and 34 sites in adjacent, downstream Grand Canyon National Park (spanning ∼464 km of river) with environmental DNA and traditional surveys. However, we did not detect salamanders elsewhere, matching prior information that salamanders are rare in the Colorado River corridor below Glen Canyon Dam. Based on this information, we discuss management options for the local population of Arizona Tiger Salamanders.

## Introduction

Conservation is often complicated. For example, actions on lands managed by the U.S. National Park Service (NPS) must consider ecological, cultural, and historical perspectives or outcomes, and some actions may seem to conflict with the Organic Act that established the agency (16 U.S.C. § 1 1916). In the southwestern USA, the Colorado River is a particularly complicated case in point, with multiple stakeholders and parties invested in management and conservation outcomes that are tied to shrinking river flows and reservoir storage (Schmidt et al. 2023).

The NPS is evaluating proposed management actions to limit establishment and impacts from non-native predators on rare native fishes and other aquatic species in the Colorado River, Arizona (NPS 2024). Since Glen Canyon Dam was completed in 1963, cold hypolimnetic releases from Lake Powell limited establishment of non-native, warmwater fishes in the Colorado River immediately downstream of the dam. However, several decades of drought in the region have caused abnormally low reservoir levels in Lake Powell. During 2022–2023, these low waters facilitated passage through, and reproduction below, Glen Canyon Dam by Smallmouth Bass (*Micropterus dolomieu*) and other non-native fishes (Eppehimer et al. 2024).

The warm waters (>15.5°C) are unusual in the modern Colorado River and enabled local spawning and downstream spread of Smallmouth Bass, causing concern they will prey upon and threaten the persistence of the federally threatened Humpback Chub (*Gila cypha*) and federally threatened Razorback Sucker (*Xyrauchen texanus*), as well as other fishes endemic to the Colorado River basin, including Flannelmouth Sucker (*Catostomus latipinnis*) (NPS 2024).

One of the primary spawning and rearing areas for Smallmouth Bass and Green Sunfish is a slough (informally called river mile -12 slough [hereafter, -12 mile slough]; ∼2.5 ha; Table 1), approximately 4.8 km below Glen Canyon Dam (NPS 2024). This slough has slow-flowing or standing water and abundant aquatic macrophytes, which are uncommon habitat features in these canyon bottomlands (NPS 2024). Green Sunfish (*Lepomis cyanellus*) were first documented breeding in -12 mile slough in 2015, whereas Smallmouth Bass were first detected spawning in the slough in 2022 (NPS 2024). Smallmouth Bass, which are a larger management concern than Green Sunfish, have continued to spawn in -12 mile slough and have increased in abundance in the approximately 32-km stretch of river downstream of the dam since 2022 (Shollenberger 2024).

**Table 1:** Amphibian species detected via either environmental DNA or visual and dip-net surveys in Glen Canyon National Recreation Area (Arizona, USA), by sampling date. Environmental DNA filters were only analyzed for Western Tiger Salamanders (AMMA), Northern Leopard Frogs, Red-spotted Toads, and Woodhouse’s Toads (ANWO).

| Site | Latitude | Longitude | 18 April 2024 | 30–31 July 2024 |
| --- | --- | --- | --- | --- |
| hidden slough | 36.8744 | -111.5565 | ANWO | ANWO |
| Horseshoe Bend marsh | 36.8771 | -111.5163 | ANWO | ANWO |
| river mile -12 slough (upper) | 36.8999 | -111.5055 | AMMA, ANWO | AMMA, ANWO |
| dam slough | 36.9268 | -111.4806 | no species detected | no species detected |

Anticipating a future in which low-water conditions like those during 2022–2023 are more common (Eppehimer et al. 2024), the NPS is considering a management option to channelize -12 mile slough so it is part of the river main stem, with the goal of increasing flow velocity, reducing water temperatures, and ultimately reducing reproduction and abundance of Smallmouth Bass and other non-native fishes (NPS 2024). Notably, -12 mile slough is one of only four sloughs or backwaters in the 24-km stretch of river within Glen Canyon National Recreation Area (NRA) below Glen Canyon Dam. The upper end of this slough is spring fed, has water year-round, and loses surface water connection with the lower parts of the slough and Colorado River seasonally (NPS 2024). The locally rare, spring-fed habitat present in the upper portion of the slough also provides habitat for other species in Glen Canyon NRA, including a population of Western Tiger Salamanders (*Ambystoma mavortium* subsp.) (NPS 2024).

To provide information for pending management decisions, we sought to (1) determine the taxonomic identity of salamanders present in -12 mile slough; (2) characterize the amphibian community in upper portion of -12 mile slough and the other three sloughs or backwaters below the dam in Glen Canyon NRA; and (3) provide information on the broader distribution of tiger salamanders and other amphibians within the Colorado River corridor below Glen Canyon dam. Determining the taxonomic identity of the salamanders was important because of uncertainty regarding whether they are native or were introduced, possibly via the bait trade (e.g., Jones et al. 1988). To accomplish objectives two and three, we used visual surveys and environmental DNA (eDNA) surveys of the four sloughs in Glen Canyon NRA below the dam during spring and summer 2024 and incorporated eDNA samples collected during summer 2020 from 34 sites along or near the Colorado River within Grand Canyon National Park (NP). Collectively, these samples span ∼464 km of river from Glen Canyon NRA through Grand Canyon NP and represent most known sloughs, backwaters, and springs within the river canyon.

## Materials and Methods

### Study area and species

Surveys for amphibians in Glen Canyon NRA below Glen Canyon Dam included trapping sloughs with baited minnow traps, visual and dip-net surveys, and eDNA surveys (Fig. 1). We use the informal names for these sites: hidden slough, Horseshoe Bend marsh (called leopard frog marsh in NPS [2004]), -12 mile slough, and dam slough, following usage in NPS (2024) and Drost et al. (2011) (Table 1). The eDNA analyses targeted only a subset of the amphibian community (Persons and Nowak 2008; Holycross et al. 2022), but included local priority species (Western Tiger Salamander, Northern Leopard Frog [*Lithobates pipiens*]) and species we expected were common but their larvae can be difficult to distinguish morphologically (Red-spotted Toad [*Anaxyrus punctatus*], Woodhouse’s Toad [*A. woodhousii*]). While none of the four species we targeted are federally listed, the Northern Leopard Frog is a species of conservation concern because of widespread declines in the region (Clarkson and Rorabaugh 1989; Drost et al. 2011).

**Figure 1.**
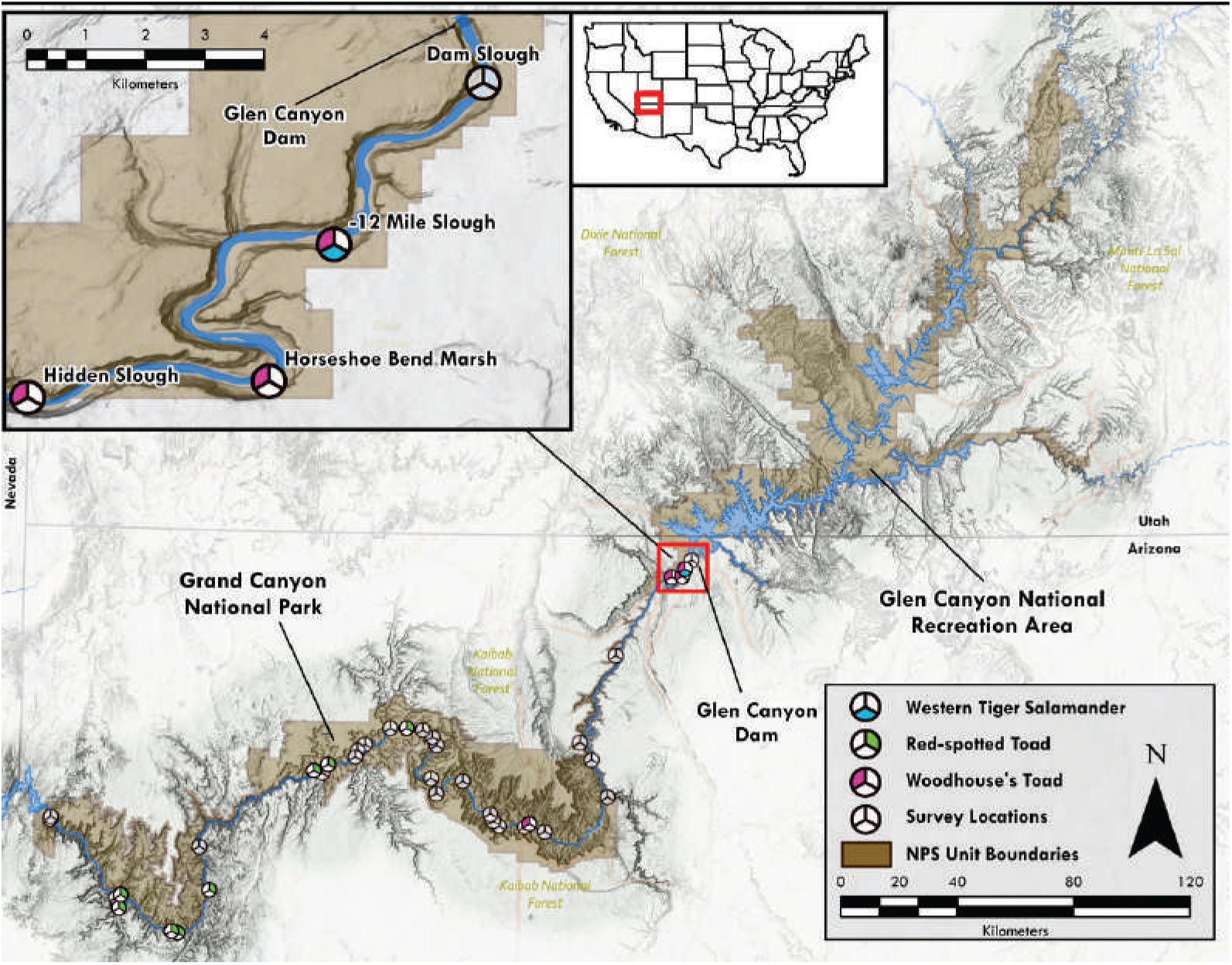
Distribution of sampling effort for amphibians in Glen Canyon National Recreation Area (inset, top left) and Grand Canyon National Park (Arizona, USA). Filled wedges indicate detection of species via either environmental DNA (eDNA) or day-time visual and dip-net surveys during 2024. Sites in Grand Canyon National Park were surveyed via eDNA only. (Data sources for map: ESRI, National Park Service, US Census Bureau, USGS, USDA Forest Service, Bureau of Transportation Statistics, Federal Highway Administration, Transport Canada, Natural Resources Canada, and the Mexican Transportation Institute.)

### Taxonomic identification, Glen Canyon National Recreation Area

The Western Tiger Salamander is comprised of several sub-species or lineages across western North America (Shaffer and McKnight 1996). Because the sub-species Barred Tiger Salamander (*A. mavortium mavortium*) was moved widely as part of the bait trade and became established in many areas in the Southwest, Western Tiger Salamander populations often represent native and non-native genotypes (Jones et al. 1988; Johnson et al. 2011). The various sub-species cannot always be distinguished visually, especially hybrids. To determine if the salamanders in -12 mile slough were the native genotype (Arizona Tiger Salamander [*A. mavortium nebulosum*]) or non-native genotypes, we used tissue samples collected from nine post-metamorphic salamanders captured in minnow traps during spring 2024. Park staff also trapped the other sloughs but did not capture any salamanders. We collected a toe-or tail-tip from each salamander and stored it in 100% ethanol until DNA extraction to determine genetic identification of the salamanders. These nine tissue samples were compared to ten tissue samples collected by researchers from Northern Arizona University during 2019 on the Kaibab Plateau in northern Arizona as well as samples previously collected from that area that we were confident represented the native genotype of salamanders (Johnson et al. 2011). The 2019 tissue samples were collected from larvae inhabiting two locations (five from Fracas Lake: 36.6306, -112.2386 and five from an unnamed stock pond [Joe’s mudhole #1]: 36.5768, -112.2117) and stored in a laboratory freezer (≤ -20°C) until analyses.

We extracted DNA from these 19 tissue samples using a DNeasy Blood & Tissue Kit (Qiagen, Inc. [Hilden, Germany]). We used primers THR and 651 from Shaffer and McKnight (1996) to amplify a 1192 base pair portion of the mitochondria (D-loop and adjacent insert) and submitted the resulting PCR product to Arizona Research Labs (Tucson, USA) for Sanger sequencing. We then aligned and analyzed these sequences in Sequencher (Gene Codes Corporation [Michigan, USA]).

### Visual and eDNA surveys, Glen Canyon National Recreation Area

Surveys of hidden slough, Horseshoe Bend marsh, the upper portion of -12 mile slough, and dam slough in Glen Canyon NRA occurred on 18 April 2024 and all sites were sampled again on 30–31 July 2024 (Table 1). We did not survey the lower portion of -12 mile slough because consistent trapping there since 2022 had not produced any captures of tiger salamanders (J. Arnold, pers. obs.), our primary target during eDNA surveys during 2024. Visual surveys were conducted during daylight hours and included scanning the water column and shoreline for animals and sweeping vegetated areas with dip nets (Hossack et al. 2022).

We collected two to four eDNA samples (volume: 1.0 – 10.4 L/filter) from each of the four sloughs in Glen Canyon NRA (Table 1). Each sample was collected from a short segment (e.g., < 30 m) of shoreline that represented suitable habitat for local amphibians. We filtered water directly from sloughs using clean techniques with a peristaltic pump connected to 5-µm pore size polyethersulfone filter membrane in a self-preserving eDNA filter pack (Smith-Root [Washington, USA]). Once the filter was nearly clogged, we dried it by pumping air through the housing for 1 minute, left the filter in its housing, and stored each filter in its original pouch with added silica beads under cool (≤ 20°C), dry conditions. We shipped filters in August 2024 to Washington State University, where we extracted DNA using the Qiashredder/DNeasy method described in Goldberg et al. (2011). Filter extractions and qPCR set up were performed in a laboratory dedicated to low-quantity DNA samples, where no tissue samples have been handled, and researchers are required to shower and change clothes before entering after being in a high-quality DNA or post-PCR laboratory.

All samples were analyzed in triplicate using assays for Western Tiger Salamanders (Moss et al. 2022), Northern Leopard Frogs (Randall et al. 2023), Red-spotted Toads (eastern and western clade assays; Appendix 1), and Woodhouse’s Toads (Appendix 1). The Western Tiger Salamander assay does not distinguish among sub-species and therefore does not provide information on whether salamanders, if detected from eDNA surveys, were native or non-native genotypes. We included a negative extraction control with each set of extractions and an additional negative qPCR control with each plate of samples.

We included an internal positive control (Qiagen [Hilden, Germany] or Thermo Fisher Scientific [Massachusetts, USA]) in each well to test for PCR inhibition. If there was evidence of inhibition, samples were cleaned with a OneStep™PCR Inhibitor Removal Kit (Zymo [California, USA]) and re-run. We considered a positive sample as any with exponential amplification in all three wells after it was first tested, or ≥ 0.33 from two triplicate reactions (samples were re-run in triplicate if the original triplicate wells yielded inconsistent results). We included a four-level standard curve on each plate consisting of a DNA sample extracted from tissue samples for each respective taxon, diluted 10^-3^ through 10^-6^ in dilution buffer and run in duplicate.

### eDNA surveys, Grand Canyon National Park

Samples for eDNA were collected from 34 sites in Grand Canyon NP during 19–27 August 2020 as part of a different project assessing distributions of native and non-native, wetland-dependent taxa. These samples were originally submitted to U.S. Forest Service National Genomics Center for Wildlife and Fish Conservation (Missoula, Montana, USA). Field samples of water were filtered with 1.5-μm, glass fiber microfilters, with the goal of sampling 5 L per site. At 29 sites, 5 L was filtered using only one filter, whereas three sites required two filters to process 5 L, and only 1.5 L was filtered (one filter) at a single site. Filters were stored in clean sample bags with silica beads until shipment to the laboratory, where DNA was extracted following standardized methods (see Franklin et al. 2019).

Extracted DNA from these 34 sites was originally analyzed for several species or pathogens. The remaining extract volume was stored in a laboratory freezer at ≤ -20°C at the U.S. Forest Service eDNA laboratory. Because that laboratory did not have a validated protocol for the Western Tiger Salamander, we shipped the extracts during September 2024 to the Washington State University laboratory and analyzed them for the same four species as from the Glen Canyon NRA samples using the methods described above.

## Results

### Taxonomic identification

We found two distinct haplotypes of tiger salamanders from the upper portion of -12 mile slough in Glen Canyon NRA, both identical to sequences documented from native populations elsewhere in Arizona. Six of the tissue samples from -12 mile slough were identical to a sequence from the 2019 samples from Fracas Lake, Arizona (Kaibab Plateau), approximately 75 km to the southwest. This Fracas Lake sample had a 1-base pair change from the other 2019 samples from that site, which were perfect sequence matches to a historical sample from that same site (HBS7832; Genbank accession number HM544138.1; Johnson et al. 2011). The three other samples from -12 mile slough had sequences identical to historical sample HBS1467 collected from Clint’s Well, Arizona (34.454 -111.396; Genbank accession number HM544137.1; Johnson et al. 2011), approximately 270 km south of -12 mile slough.

### Visual and eDNA surveys, Glen Canyon NRA

We detected two species of amphibians in Glen Canyon NRA below the Glen Canyon Dam (Table 1, Fig. 1). Salamander DNA was detected only from the upper portion of -12 mile slough. All four filters collected from the upper portion of the slough on 18 April 2024 tested positive for presence of Western Tiger Salamanders, but only one of four filters collected on 31 July 2024 was positive. No salamanders were detected via visual or dip-net surveys in the upper portions of the slough, which is not surprising because those methods have low detection probabilities unless animals are abundant, especially when there is extensive cover or presence of fish that can cause salamanders to hide (Hossack et al. 2022).

The Woodhouse’s Toad was detected from three of four sites in April and July 2024 (Table 1). Toad tadpoles too small to be identified in the field were seen at hidden slough and Horseshoe Bend marsh on 18 April 2024. The eDNA samples from both sites tested positive for the Woodhouse’s Toad and not the Red-spotted Toad, suggesting the tadpoles were the former species. No amphibians were detected at dam slough during April or July 2024 surveys and the Northern Leopard Frog was not detected at any sites.

### eDNA surveys, Grand Canyon NP

Tiger salamanders and Northern Leopard Frogs were not detected at any of the 34 sites sampled via eDNA in Grand Canyon NP. Red-spotted Toads were detected at nine sites and Woodhouse’s Toads were detected at one site. Red-spotted Toads were detected more frequently as sampling proceeded west (Fig. 1).

## Discussion

Our analysis of tissue samples from nine metamorphosed salamanders from -12 mile slough indicates they are native Arizona Tiger Salamanders (*Ambystoma mavortium nebulosum*). We found sequence matches with a population from the Kaibab Plateau in northwestern Arizona (∼75 km away) and a more distant population on the Mogollon Plateau in central Arizona (∼270-km south). The upper portion of -12 mile slough was also the only site where salamanders were detected via eDNA sampling from four sites in Glen Canyon NRA below the dam and 34 sites in Grand Canyon NP. Our analysis of tissue samples was based only on mitochondrial DNA, so we cannot rule out past hybridization with non-native Barred Tiger Salamanders, but if that were the case, the maternal lineage that survived was from native Arizona Tiger Salamanders.

There has been uncertainty regarding the origin of the salamanders in -12 mile slough since they were first documented in 2016 (NPS 2024). The local population might be a relict, like the Northern Leopard Frog population that used to inhabit the same stretch of river (Drost et al. 2011). Although salamander populations inhabiting riverine habitats of the inner canyon below Glen Canyon Dam or in the Grand Canyon seem rare (Miller et al. 1982; Albright et al. 2022), Arizona Tiger Salamanders were collected from multiple tributaries, including the Navajo Creek watershed, and near the Colorado River in 1938, upstream of the eventual dam location (Woodbury 1959). A tiger salamander was seen at Horseshoe Bend Marsh in Glen Canyon NRA in 2015 (NPS 2024), but we are unaware of other records outside of -12 mile slough indicating a resident population downstream of the dam in Glen Canyon NRA (Albright et al. 2022). However, spring-fed marsh habitats are also rare along the river corridor, so there may be few locations suitable to support salamanders. For example, of the 34 sites sampled in Grand Canyon NP during 2020, only one had marsh habitat resembling that of -12 mile slough (B. Holton, NPS, pers. comm., 07 Nov 2024).

If salamanders did not colonize -12 mile slough naturally, there are several plausible ways they could have been introduced. Tiger salamander larvae were commonly used for fishing bait in this region, including for use at Lake Powell (https://home.nps.gov/glca/learn/nature/amphibians.htm), although salamanders are no longer allowed as bait in Utah or in Glen Canyon NRA (Utah administrative rule R657-13-12; https://www.nps.gov/glca/planyourvisit/fishing.htm). Our analysis of mitochondrial DNA indicates the population in -12 mile slough is not comprised of Barred Tiger Salamanders, but it is possible the population was founded by salamanders collected as bait from local sources. For example, Arizona Tiger Salamanders are common in waterbodies on the Kaibab Plateau ∼60 km southwest, where our reference samples for mitochondrial DNA originated. We found matching sequences for a population from the Kaibab Plateau (Fracas Lake) for six of nine salamanders sampled from -12 mile slough. But 3 salamanders had matching sequences from a population of Arizona Tiger Salamanders in central Arizona. These two groups of Arizona Tiger Salamander sequences, 11 base pairs apart, suggest two origins for this population, for example a secondary influx of migrants (or introductions) into the area or migrants (or introductions) from a mixed source.

Tiger salamanders were also present in ponds on the Lake Powell National Golf Course in Page, Arizona, which is 3.5-km upstream of -12 mile slough. These ponds were 800 m from the canyon rim and were being evaluated for rearing Razorback Suckers during the late 1990s (Mueller and Wick 1998). A large flood during September 2013 caused municipal ponds in the same area to overfill and water was reported flowing into the canyon upstream of -12 mile slough (Wildman Jr and Vernieu 2017), suggesting the floods may have aided colonization by salamanders down to the canyon. We were unaware of this potential local salamander population in Page when we gathered tissue samples for analysis and therefore cannot evaluate if that population could have been the source of salamanders -12 mile slough. We also cannot determine whether salamanders colonized ponds in Page naturally or were introduced intentionally. But if they were inadvertently transported with Razorback Suckers from the Ouray National Fish Hatchery located in Vernal, Utah, to these ponds in Page (e.g., Green and Dodd 2007) and subsequently became established in the river canyon, we would have expected sequence matches with salamanders from the Vernal, Utah, area. Instead, tiger salamander sequences from the Vernal area fall within the Great Plains clade of *A. mavortium nebulosum* (R46 in Johnson et al. 2011), rather than Southwest clade of *A. mavortium nebulosum* (R05 and R08 in Johnson et al. 2011). Notably, a citizen science observation from May 2024 (https://www.inaturalist.org/observations/217387859) suggests tiger salamanders are still in Page. Comparing DNA sequences of salamanders from -12 mile slough, Page, and elsewhere in the surrounding area could help resolve the origin of salamanders in and around Glen Canyon NRA.

The upper portion of -12 mile slough, with both tiger salamanders and Woodhouse’s Toads documented via eDNA, was also the only site in either Glen Canyon NRA or Grand Canyon NP where we detected more than 1 species of amphibian. Overall, we detected Woodhouse’s Toads at three of four sites in Glen Canyon NRA below the dam but at only one of 34 sites in Grand Canyon NP. In contrast, we did not detect Red-spotted Toads in Glen Canyon NRA but detected them at nine sites in Grand Canyon NP. We are unsure why we detected these species so rarely since both occur widely in the area and other recent surveys have indicated they are abundant (e.g., Persons and Nowak 2008; Corsetti and Holton 2020, Albright et al. 2022). We suspect the lack of nighttime surveys reduced detection of juvenile and adult toads (e.g., Bradford et al. 2005), and the late-summer eDNA sampling of Grand Canyon may have occurred after most larvae metamorphosed and departed breeding sites (e.g., Halstead et al. 2023), making them unavailable for detection.

Based on eDNA samples, we also detected Red-spotted Toads only west of the approximate split between the eastern and western clades of this species (Jaeger et al. 2005). This suggests there could be undocumented sequence variation to the east that reduced specificity of eDNA primers. However, the eastern clade assay was a perfect match for a previous tissue sample collected from the junction with the Colorado and Little Colorado rivers (LVT6051; haplotype E01; GenBank accession number DQ085757.1; Jaeger et al. 2005). Regardless, we recommend future surveys intended to evaluate occurrence of amphibians in fringe wetlands, backwaters, and springs along the Colorado River corridor use multiple survey methods (e.g., VES and eDNA) timed to capture early season (April–May) and late-summer, post-monsoon breeding activity (Corsetti and Holton 2020; Holycross et al. 2022), and collect Red-spotted Toad tissues for further validation of eDNA primers.

Our surveys were not intensive and only occurred during daytime, many potential species were not seen or tested for presence via eDNA, and our inferences are not grounded in model-based estimates that account for detection uncertainty (e.g., Moss et al. 2022; Hossack et al. 2023). Therefore, lack of detections from specific sites should not be considered definitive. We relied on eDNA because of time constraints and because it has proven more effective for tiger salamanders in the region compared to visual and dipnet surveys (Hossack et al. 2022), although detection of tiger salamanders via eDNA may still be lower than for other co-occurring amphibian species in the region, especially in stream-associated habitats or after arrival of summer rains that can induce dispersal away from aquatic habitats (Hossack et al. 2023).

Locally, adult Western Tiger Salamanders typically hibernate underground during the winter, migrate to waterbodies during early spring to breed and then return to terrestrial environment, and larvae metamorphose and disperse into the terrestrial environment after summer rains (Collins 1981, Brunner et al. 2004). These life history characteristics and their timing can constrain detection of aquatic animals and may be important for potential mitigation strategies, discussed below.

The lack of detections of Northern Leopard Frogs from any of our surveys likely reflects the species’ widespread decline in the region, especially near historical range margins such as northern Arizona (Clarkson and Rorabaugh 1989; Drost et al. 2011). As of 2011, all previously known Northern Leopard Frog populations along the Colorado River in Arizona were considered extirpated (Drost et al. 2011). Horseshoe Bend Marsh in Glen Canyon NRA is the last place in Glen Canyon NRA where Northern Leopard Frogs were known to occur, with the population disappearing around 2005 (called Horseshoe Bend or -9 mile draw in Drost et al. 2011). That site became densely vegetated with common reeds (*Phragmites australis*) and has abundant Green Sunfish (J. Arnold, pers. obs.). In Grand Canyon NP, the last river-associated habitat known to support Northern Leopard Frogs apparently no longer holds water (called cardenas marsh in Drost et al. 2011).

Potentially converting a rare, spring-fed wetland into flowing, riverine habitat to slow downstream invasion by non-native predators that threaten rare fishes illustrates the challenges and complexity of conservation in this system. If -12 mile slough is channelized, our data and prior data indicating salamanders occur only or primarily at that site within the river canyon, suggests they may be functionally extirpated from the river corridor in Glen Canyon NRA. If the timing is appropriate and salamanders hibernate away from -12 mile slough (e.g., near rock walls), capturing salamanders as they migrate to the slough and relocating them to a habitat nearby is a potential management option, given evidence that the resident salamanders are native to the region. There are limited data on success rates of relocating salamanders, however, especially with a presumably small number of individuals available from a source population (e.g., Germano and Bishop 2009). Whether relocating salamanders is a preferred management option would likely depend on whether they are suspected to have colonized the canyon naturally.

## Acknowledgments

We thank Mary Sterling for processing the eDNA and tissue samples at Washington State University, Bayley Stevens for assistance in the field, and Joe Mihaljevic and Kelsey Banister (Northern Arizona University) for sharing 2019 tissue samples. We also thank Brandon Holton (NPS) for sharing his eDNA samples and providing feedback on the text and the U.S. Forest Service National Genomics Center for Wildlife and Fish Conservation for providing archived samples. Comments from Charles Drost and Charles Yackulic improved the manuscript. Work was done through Arizona Game and Fish Department scientific collection license SP650682 to Joe Mihaljevic (NAU IACUC protocol 18-010), Arizona Game and Fish Department scientific collection license SP648297 to Blake Hossack (UM IACUC protocol 065-23BHWB-012924), and the 2018 Expanded Non-native Aquatic Species Management Plan in Glen Canyon National Recreation Area and Grand Canyon National Park below Glen Canyon Dam (https://parkplanning.nps.gov/projectHome.cfm?projectId=74515). This is Amphibian Research and Monitoring Initiative (ARMI) product number ZZZ. Any use of trade, firm, or product names is for descriptive purposes only and does not imply endorsement by the U.S. Government.

Work was done through Arizona Game and Fish Department scientific collection license SP650682 to Joe Mihaljevic (NAU IACUC protocol 18-010), Arizona Game and Fish Department scientific collection license SP648297 to Blake Hossack (UM IACUC protocol 065-23BHWB-012924), and the 2018 Expanded Non-native Aquatic Species Management Plan in Glen Canyon National Recreation Area and Grand Canyon National Park below Glen Canyon Dam (https://parkplanning.nps.gov/projectHome.cfm?projectId=74515). Funding for assay development and validation was provided under cooperative agreement P22AC00993-01 from the National Park Service Inventory and Monitoring Program. C. Goldberg was supported in part by the USDA National Institute of Food and Agriculture, McIntire-Stennis project 1018967. Any use of trade, firm, or product names is for descriptive purposes only and does not imply endorsement by the U.S. Government.

## Appendix

### eDNA assay design and validation for Red-spotted Toad (*Anaxyrus punctatus*)

We designed this environmental DNA (eDNA) assay using cytochrome b sequences from Jaeger et al. (2005) and Bryson et al. (2012). There was too much variation among the clades (western, eastern, and peninsular) for a single assay, so we developed a separate assay for each of the western and eastern clades and left out the peninsular clade; therefore, this assay should not be used for samples collected on the Baja California peninsula (Mexico). Also, results from this study suggest there may be undocumented haplotype variation in the upper Grand Canyon area that could produce false negatives, so we recommend developing sequencing data from tissue samples in that area to check for assay mismatches.

We developed an inclusive consensus sequence for each of the clades (western, eastern) and used Primer Express (Thermo Fisher Scientific) to produce designs for quantitative PCR (qPCR) assays. We then validated those designs in silico using Primer-BLAST (Ye et al. 2012) and ordered primers and a probe for each of the clades that were not predicted to amplify co-occurring species (Table A1). We note that the assay for the eastern clade (BUPU-E) may amplify other frogs and fish not found in the range of this species, as well as Blue Catfish (*Ictalurus furcatus*), which are not present in the Colorado River upstream of Lake Mead (Table A1). We validated those assays in multiplex (eastern, western, and internal control; Qiagen, Inc.) using tissue samples (Table A2). The total solution volume was 15 µl and consisted of 1X Environmental Master Mix (Thermo Fisher Scientific), 0.2 µm concentration of each primer and probe, and 3 µl of DNA extract. The cycling protocol started with 10 minutes at 95°C followed by 50 cycles of 95°C for 15 seconds and 60°C for 60 seconds. Reactions were run on a CFX96 Touch Real-Time PCR Detection System (Bio-Rad).

**Table A1:**
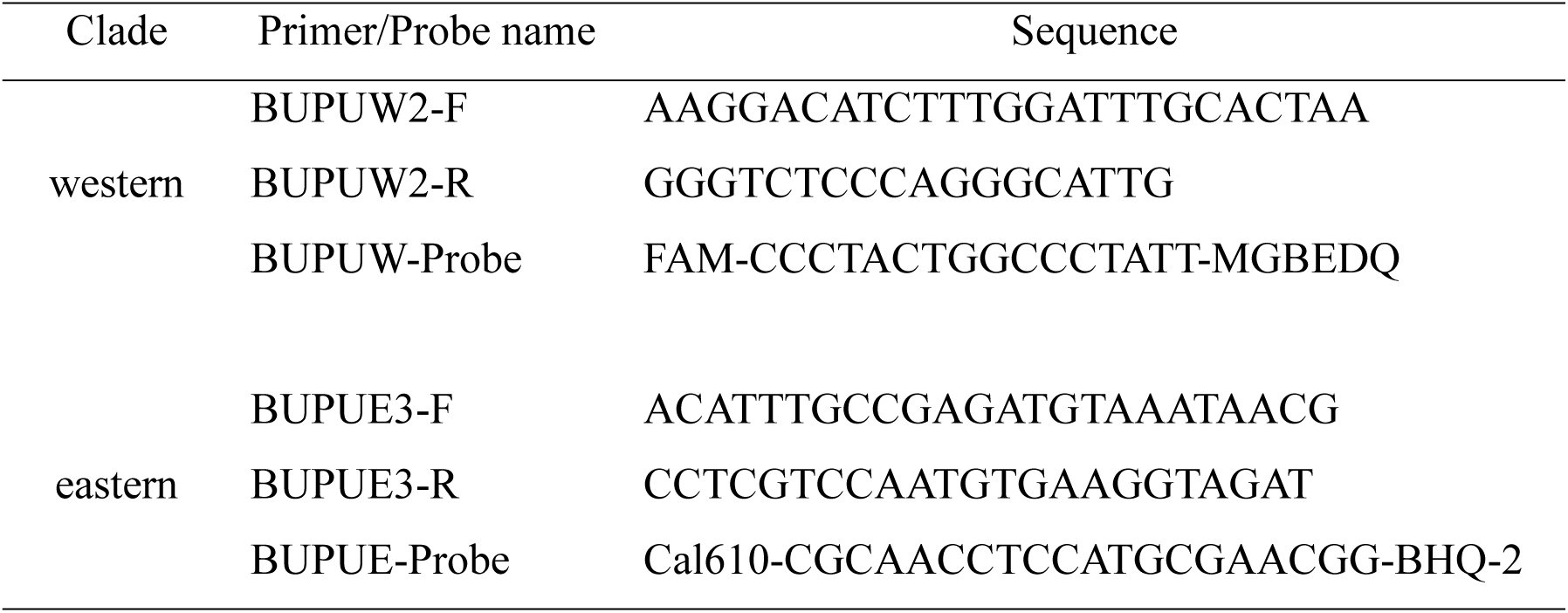
Primers and probes for environmental DNA (eDNA) assays for the western and eastern clades of the Red-Spotted Toad (*Anaxyrus punctatus*).

**Table A2:** Tissue samples used to validate environmental DNA (eDNA) assays for the Red-spotted Toad (*Anaxyrus punctatus*) and Woodhouse’s Toad (*Anaxyrus woodhousii*).

| Scientific name | Common name | Number of samples |
| --- | --- | --- |
| *Anaxyrus cognatus* | Great Plains Toad | 5 |
| *Anaxyrus microscaphus* | Arizona Toad | 5 (10 for *A. woodhousii*) |
| *Anaxyrus punctatus* | Red-spotted Toad | 6 (5 for *A. woodhousii*) |
| *Anaxyrus woodhousii* | Woodhouse's Toad | 5 (14 for *A. woodhousii*) |
| *Incilius alvarius* | Sonoran Desert Toad | 5 |
| *Hyla [Dryophytes] arenicolor* | Canyon Treefrog | 5 |
| *Hyla [Dryophytes] wrightorum* | Arizona Tree Frog | 5 |
| *Pseudacris maculata* | Boreal Chorus Frog | 5 |
| *Pseudacris sierra* | Sierran Treefrog | 5 |
| *Lithobates catesbeianus* | American Bullfrog | 5 |
| *Lithobates chiricahuensis* | Chiricahua Leopard Frog | 5 |
| *Lithobates yavapaiensis* | Lowland Leopard Frog | 5 |
| *Lithobates pipiens* | Northern Leopard Frog | 5 |
| *Scaphiopus couchii* | Couch's Spadefoot | 5 |
| *Spea intermontana* | Great Basin Spadefoot | 5 |
| *Spea multiplicata* | Mexican Spadefoot | 5 |

### eDNA assay design and validation for Woodhouse’s Toad (*Anaxyrus woodhousii*)

We designed this assay using cytochrome b sequences we developed using MVZ28 and MVZ43 primers (Graybeal 1993) from five tissue samples collected in Arizona, California, and Idaho (USA). We used a gel extraction to isolate the correct PCR product from an agarose gel and sequenced this product using Sanger sequencing.

We manually designed this assay to ensure that it would not amplify the closely related Arizona Toad (*A. microscaphus*; Table A3). We evaluated resulting designs using Primer Express (Thermo Fisher Scientific) and Primer-BLAST (Ye et al. 2012) and ordered a set of primers and probe predicted to not amplify any other species (Table A3). We validated this assay (including internal control; Qiagen, Inc.) using tissue samples (Table A2), including Woodhouse’s Toad samples from Nevada, California, New Mexico, and Arizona (two from Glen Canyon National Recreation Area, Arizona). The total solution volume was 15 µl, with 1X Environmental Master Mix, 0.2 µm concentration of each primer and probe, and 3 µl of DNA extract. The cycling protocol started with 10 minutes at 95°C followed by 50 cycles of 95°C for 15 seconds and 60°C for 60 seconds.

**Table A3:**
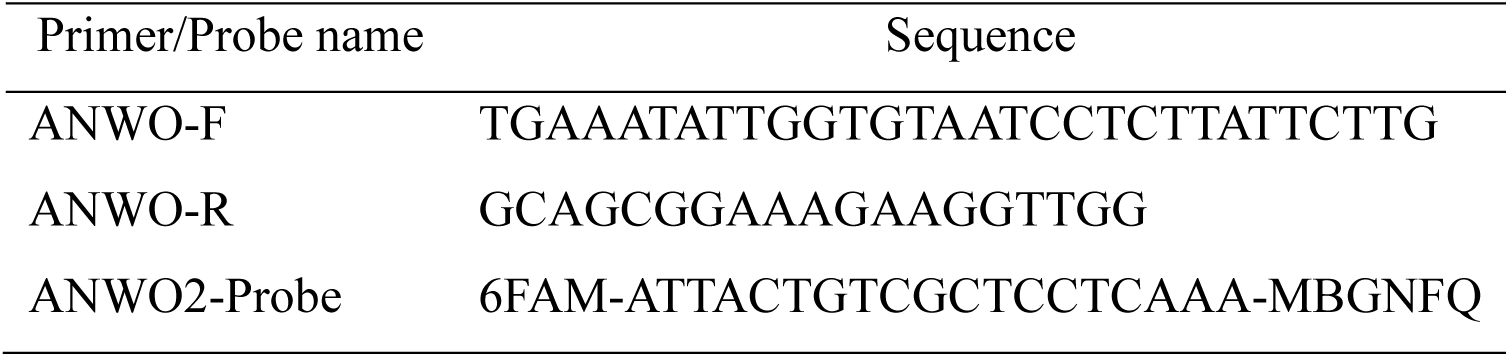
Primers and probe for the Woodhouse’s Toad (*Anaxyrus woodhousii*) environmental DNA (eDNA) assay.

